# Centralspindlin promotes *C. elegans* anchor cell specification, vulva induction and morphogenesis

**DOI:** 10.1101/2024.09.18.613761

**Authors:** Tatsuya Kato, Olga Skorobogata, Christian Rocheleau

**Affiliations:** Department of Anatomy and Cell Biology, McGill University, Montreal, Quebec, Canada; Metabolic Disorders and Complications Program, Research Institute of the McGill University Health Centre, Montreal, Quebec, Canada; Division of Endocrinology and Metabolism, Department of Medicine, McGill University, Montreal, Quebec, Canada

## Abstract

*Caenorhabditis elegans* vulval development is a relatively simple model of organ development whereby a signal from the overlying gonad induces three epithelial cells to undergo three rounds of cell division to generate 22 cells that make up the vulva. Specification of the vulva cell fates requires coordination between cell division and cell signaling via LIN-12/Notch and LET-23/EGFR pathways in the somatic gonad and the underlying epithelium. Here we characterize the positive regulation of vulval development by the centralspindlin complex, a conserved cytokinesis regulator. Centralspindlin, a heterotetramer of ZEN-4/KIF23 and CYK-4/RacGAP1, is essential for completion of cytokinesis during early embryonic cell divisions. We found that centralspindlin is required in the somatic gonad for division of somatic gonad precursor cells and hence specification of the LIN-3/EGF-secreting anchor cell critical for LET-23/EGFR-mediated vulval induction. However, the requirements for centralspindlin for cytokinesis during postembryonic development are incomplete as a binucleate anchor cell is frequently specified. The presence of the binucleate anchor cell correlates with vulva induction and demonstrates that LIN-12/Notch signaling, required for anchor cell specification, and LET-23/EGFR signaling, required for vulva induction, is largely functional in these cells. Centralspindlin is also partially required for cytokinesis of the vulval cells where it regulates vulva morphogenesis rather than induction. We also found that the GAP domain of CYK-4/RacGAP1 required for contractile ring assembly during embryonic division is not essential for vulval development. Thus, there appears to be different requirements for centralspindlin during postembryonic development of the somatic gonad and vulva as compared to early embryogenesis.

## INTRODUCTION

During development cell proliferation and cell fate specification must be coordinated to properly form functional tissues and organs. In *C. elegans*, the cell lineage is largely invariant and thus defects in cell proliferation are catastrophic for development (Sulston & Horvitz, 1977; Sulston et al., 1983). For example, loss of *zen-4* or *cyk-4* result in failure to complete cytokinesis during embryogenesis causing cells to become multinucleate and improperly differentiate (Jantsch-Plunger et al., 2000; Powers et al., 1998; Raich et al., 1998; Severson et al., 2000). ZEN-4/KIF23 dimers bind to CYK-4/RacGAP1 dimers to form a heterotetrameric complex called centralspindlin that is required for cytokinesis by bundling microtubules and promoting contractile ring assembly at the spindle midzone (Mishima et al., 2002). ZEN-4 contains a motor domain that organizes microtubule bundles (Powers et al., 1998) and a CYK-4-binding coiled coil domain (Raich et al., 1998). CYK-4 consists of a ZEN-4-binding coiled coil domain, a lipid-binding C1 domain and a GAP domain with activity toward the Rho GTPases *in vitro* (Jantsch-Plunger et al., 2000). Although there has been much debate as to the requirement and role of the GAP domain for cytokinesis, several studies support that it promotes contractile ring assembly through RHO-1/RhoA activation by binding with the RhoGEF ECT-2/ECT2 and relieving its autoinhibition (Zhang & Glotzer, 2015). Others suggest that the CYK-4 GAP domain may inactivate CED-10/Rac1 to decrease branched F-actin in the equatorial plane (Canman et al., 2008) which could make the cortex softer and more pliable for the contractile ring. There are obvious challenges with studying cytokinesis regulators during post-embryonic development; however, several studies have shown specific roles for *cyk-4* and *zen-4* in tissue morphogenesis. Centralspindlin regulates polarization of the arcade cells during early pharyngeal tubulogenesis (Portereiko et al., 2004; Von Stetina et al., 2017), while *cyk-4*, but not *zen-4*, regulates oocyte cellularization in the female germline (Lee et al., 2018). A recent study found that centralspindlin regulates morphogenesis of the spermatheca via its role in cytokinesis with the RHO-1 and CDC-42 GTPases (Zhang et al., 2023). We sought to determine the requirements for centralspindlin during vulva development.

The vulva in *C. elegans* is an excellent system to study organ development. Induction of the vulva relies on signaling between the developing somatic gonad and the underlying ventral hypodermal (epidermal) cells (Sternberg, 2005). LIN-12/Notch signaling plays a critical role during early somatic gonadogenesis in specifying the anchor cell versus ventral uterine cell fates (Fig. 1) (Greenwald et al., 1983; Yochem et al., 1988). Once specified, the anchor cell expresses the LIN-3/Epidermal Growth Factor (EGF) ligand and, via LET-23/EGF Receptor (EGFR), induces three of the six underlying hypodermal cells termed vulval precursor cells (VPCs; P3.p to P8.p) to generate vulval cells (Hill & Sternberg, 1992). High LET-23/EGFR activity levels in the most proximal P6.p VPC specifies the 1° vulval fate, while graded LET-23/EGFR signaling and lateral LIN-12/Notch signaling specify the adjacent P5.p and P7.p to become 2° vulval cells (Fig. 1) (Shin & Reiner, 2018). The three induced VPCs undergo invariant divisions to become 22 vulval cells and migrate to form the vulval invagination. The uninduced P3.p, P4.p and P8.p VPCs divide once and fuse with the hypodermis, the 3° non-vulval fate (Fig. 1). Changes to this tightly coordinated signaling network result in observable developmental defects: reduced LET-23/EGFR signaling leads to a vulvaless phenotype in which fewer than three VPCs are induced, while increased signaling causes a multivulva phenotype in which more than three VPCs to be induced.

**Figure 1.**
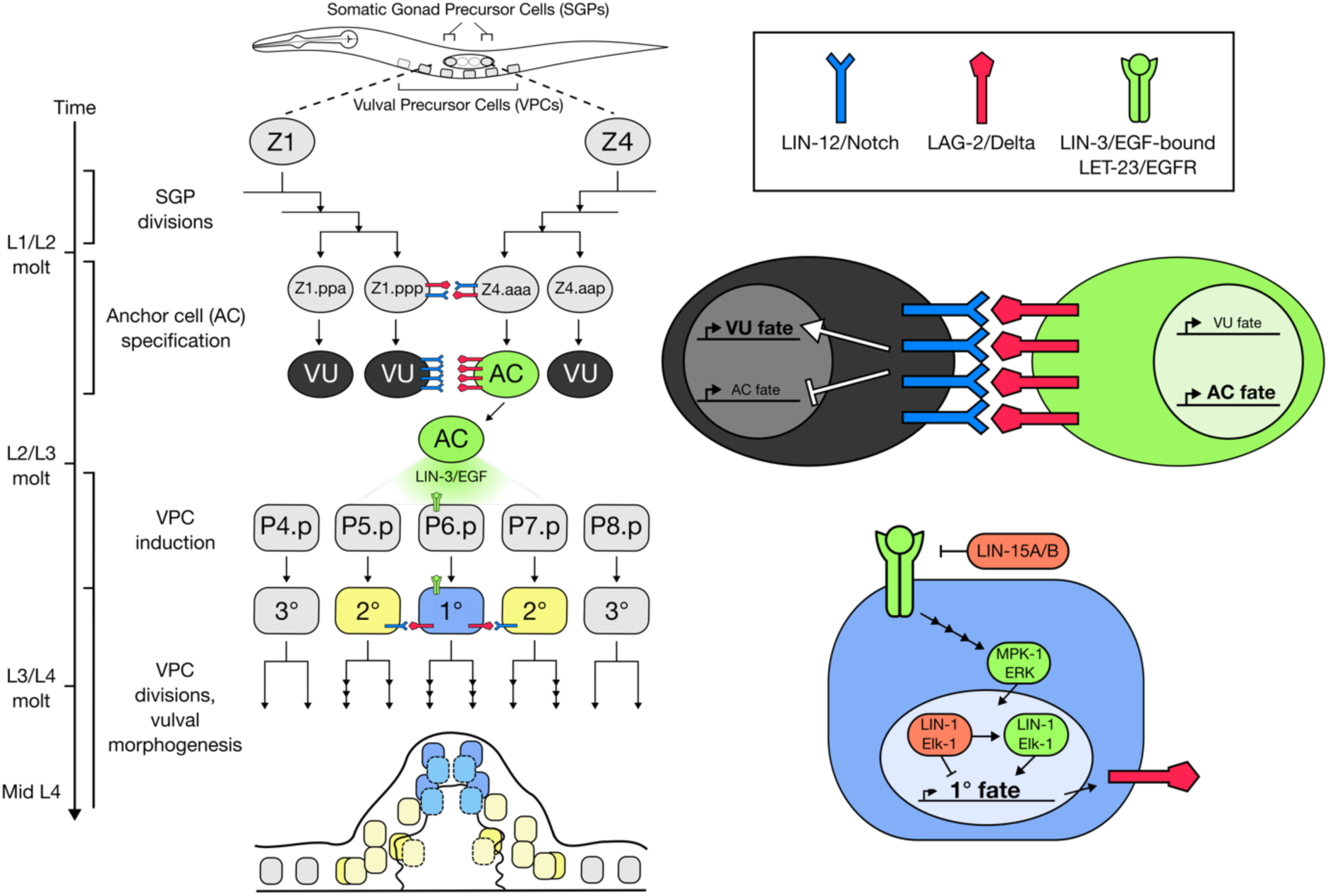
***C. elegans* vulval development requires anchor cell (AC) specification and vulval precursor cell (VPC) induction.** Left: Developmental timeline of the cell fate specification events leading to vulva formation. During the first larval stage (L1) somatic gonad precursors Z1 and Z4 divide three times to generate neighboring Z1.ppp and Z4.aaa daughters expressing LIN-12/Notch (blue) and LAG-2/Delta (red). During the second larval stage (L2) the daughter that receives the highest levels of LIN-12/Notch signal is fated to become a ventral uterine precursor (VU, black) and the other cell the anchor cell (AC, green). In early third stage (L3) larvae the AC expresses the LIN-3/EGF ligand that activates LET-23/EGFR signaling in the underlying vulva precursor cells (VPCs). P6.p receives the highest amount of signaling and becomes a primary cell (blue) and the neighboring cells P5.p and P7.p are induce by lower LET-23/EGFR signaling and later LIN-12/Notch signaling to adopt the secondary vulval fate (yellow). Right: Cartoon of LIN-12/Notch signaling during the VU/AC decision and LET-23/EGFR signaling via MPK-1/Erk during specification of the primary vulval fate. Shown in red are negative regulators of vulva induction. LIN-1/Elk1 which prevents ectopic vulva induction until activated by MPK-1/Erk and LIN-15A/B which restricts LIN-3/EGF expression to the AC. Loss of LIN-1 or LIN-15A/B results all six VPCs adopting vulval fates.

Here we demonstrate that centralspindlin is required for vulva induction and morphogenesis. We reveal that centralspindlin regulates vulva induction by controlling cytokinesis of precursor cells in the somatic gonad and hence specification of the EGF-secreting anchor cell. In animals in which a vulva has been specified, centralspindlin controls cytokinesis in the VPC daughters to regulate vulva morphogenesis. Notably, centralspindlin appears to have lower threshold requirements in the somatic gonad and the VPCs and their daughters as compared to embryonic blastomeres, suggesting that reduced requirements or redundant mechanisms contributes to cytokinesis post-embryonically.

## RESULTS

### Centralspindlin is required for vulva induction

To determine if centralspindlin regulates vulva development we primarily used *zen-4(RNAi)* and the fast-acting *zen-4(or153)* temperature sensitive allele (Severson et al., 2000). We found that *zen-4(RNAi)* starting at the L1 larval stage or shifting *zen-4(or153)* to non-permissive temperature of 26℃ at the L1 stage to avoid embryonic lethality resulted in a potent vulvaless phenotype (Fig. 2A-C). While wild type larvae had normal vulval development with symmetrical lumens (Fig. 2A), *zen-4(or153)* incubated at restrictive temperature and *zen-4(RNAi)*-treated larvae either failed to form lumens (Fig. 2B, 2C) or developed abnormally shaped lumens (Fig. 2B’). These morphologies corresponded with absent or partial VPC induction, leading to significantly fewer induced VPCs on average (Fig. 2I). The *cyk-4(t1689)* temperature sensitive coiled coil mutant (Mishima et al., 2002) as well as RNAi knockdown of *cyk-4* displayed similar vulval defects (Fig. 2D,E,I). Interestingly, the *cyk-4(or749)* temperature sensitive GAP domain mutant (Canman et al., 2008) had wild type vulval development at 26℃ (Fig. 2F,I), suggesting that the GAP activity of CYK-4 is not essential for vulval development (see discussion). Since ZEN-4 and CYK-4 could control vulval development through its role in epithelial polarity rather than regulation of cytokinesis, we determined if vulval phenotypes also arise after disruption of the cytokinesis regulators, SPD-1/PRC1 (Verbrugghe & White, 2004) and AIR-2/AURKB (Severson et al., 2000). SPD-1 is a microtubule bundler at the spindle midzone and binds to centralspindlin (Lee et al., 2015), while aurora kinase AIR-2 is a subunit of the chromosome passenger complex which promotes cortical membrane localization of centralspindlin and thus midzone contractile ring assembly (Basant et al., 2015). Loss of SPD-1 through *spd-1(oj5)* temperature shift strongly induced the vulvaless phenotype (Fig. 2G,I), while *air-2(RNAi)* knockdown disrupted vulval development similarly to *cyk-4(RNAi)* (Fig. 2H,I). Taken together, these results suggest that centralspindlin promotes vulval development likely by regulating cytokinesis.

**Figure 2.**
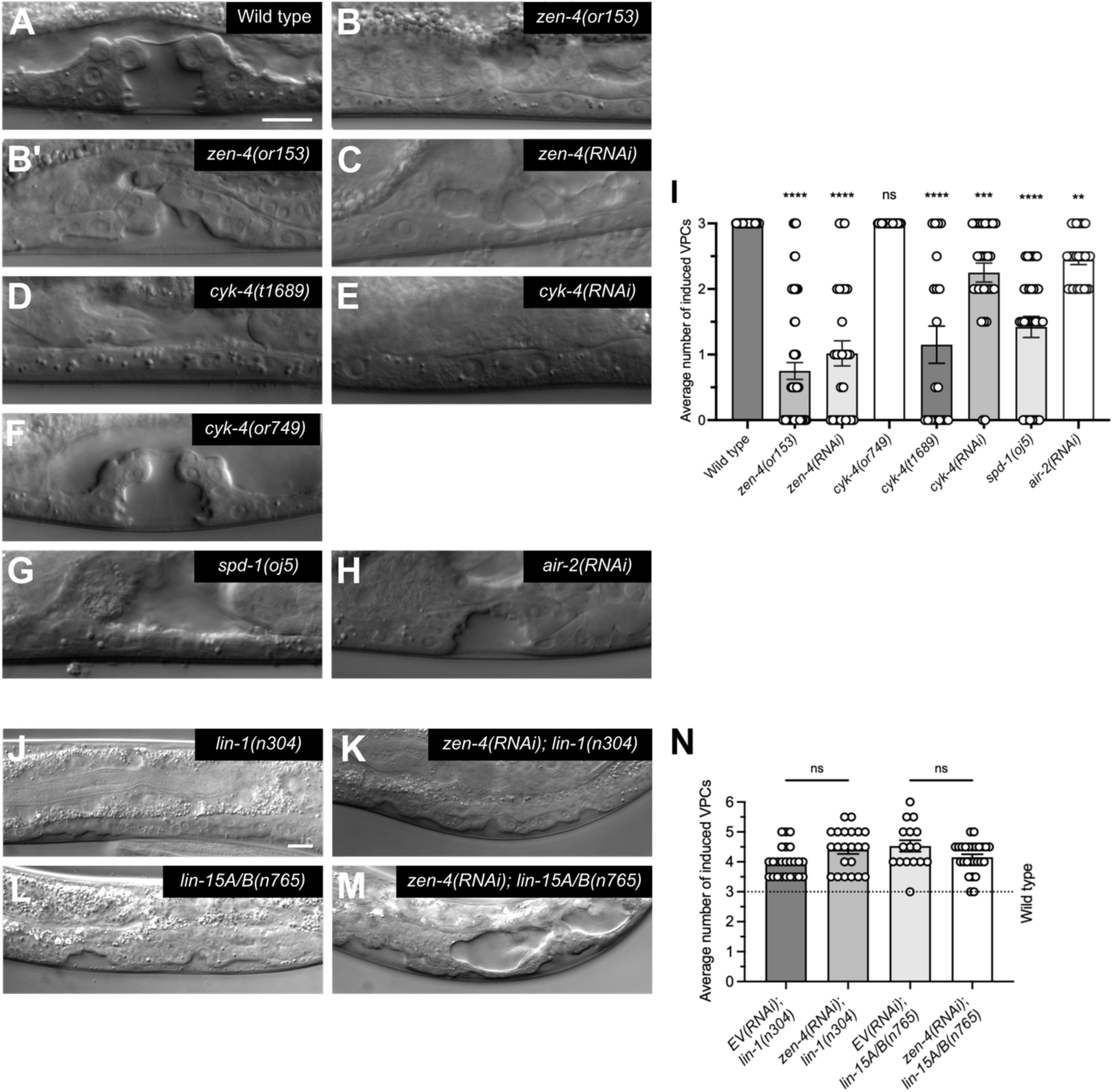
Centralspindlin promotes vulval development upstream of LET-23/EGFR signaling. (A-H) Developing vulva of wild type, *zen-4(or153)*, *zen-4(RNAi)*, *cyk-4(t1689), cyk-4(RNAi)*, *cyk-4(or749)*, *spd-1(oj5)* and *air-2(RNAi)* at mid L4. (I) VPC induction scores of wild type, *zen-4(or153)*, *zen-4(RNAi)*, *cyk-4(t1689), cyk-4(RNAi)*, *cyk-4(or749)*, *spd-1(oj5)* and *air-2(RNAi)*. Bars represent the average VPC induction scores, and dots represent individual scores. See “VPC induction scoring” in Materials and methods for details. One-way ANOVA. ns, not significant, \*\**P* < 0.01, \*\*\**P*<0.001, \*\*\*\**P*<0.0001. (J-M) Developing vulva at mid L4 stage of *lin-1(n304)* and *lin-15A/B(n765)* mutants treated with empty vector (EV) control RNAi or *zen-4(RNAi)*. (N) VPC induction scores of *lin-1(n304)*, *zen-4(RNAi); lin-1(n304)*, *lin-15A/B(n765)* and *zen-4(RNAi); lin-15A/B(n765)*. One-way ANOVA. ns, not significant. Scale bars, 10µm.

### Centralspindlin functions upstream of LET-23/EGFR signaling to promote vulva induction

We hypothesized that centralspindlin promotes vulval development by regulating VPC divisions downstream of LET-23/EGFR signaling. To determine centralspindlin’s position relative to LET-23/EGFR signaling in the VPCs, we conducted genetic epistasis of *zen-4(RNAi)* with mutants in two negative regulators of LET-23/EGFR signaling that bookend the pathway, LIN-1/ELK1 and LIN-15A/B (Fig. 1). LIN-1 is a conserved ETS-domain transcription factor and primary VPC fate inhibitor of vulva induction (Beitel et al., 1995) while LIN-15A/B are synMuv proteins which inhibit ectopic *lin-3/EGF* expression (Clark et al., 1994; Cui et al., 2006; Huang et al., 1994). Consequently, *lin-1(n304)* and *lin-15A/B(n765)* loss-of-function mutants exhibited multivulva phenotypes with ectopic lumen formations (Ferguson & Horvitz, 1985; Horvitz & Sulston, 1980) (Fig. 2J,L). Intriguingly, *zen-4(RNAi)* treatment of *lin-1(n304)* or *lin-15A/B(n765)* larvae did not suppress either multivulva phenotype (Fig. 2K,M,N), suggesting that centralspindlin is not required for the competency of the VPCs to be induced but likely functions upstream of LET-23/EGFR signaling.

### Centralspindlin promotes anchor cell specification via regulation of SGP cytokinesis

The complete absence of VPC induction in ∼50% of *zen-4(or153)* larvae (28/60; Fig. 2H), failure of *zen-4(RNAi)* to suppress the multivulva phenotypes of LET-23/EGFR negative regulator mutants (Fig. 2M) and ZEN-4’s expression in somatic gonads (Fig. S1) led us to test if centralspindlin controls vulval development by regulating the division of cells required for anchor cell specification. At the late L1 stage, two granddaughters of the Z1 and Z4 somatic gonad primordial cells called Z1.pp (i.e., the most posterior Z1 granddaughter) and Z4.aa (i.e., the most anterior Z4 granddaughter) divide (Fig. 1). Although the resulting four cells are initially competent to become either the anchor cell or a ventral uterine cell (Seydoux et al., 1990), the outer cells Z1.ppa and Z4.aap (β cells) lose competence and adopt ventral uterine cell fates while Z1.ppp and Z4.aaa (α cells) remain bipotent (Kimble, 1981; Seydoux & Greenwald, 1989; Seydoux et al., 1990). By mid L2, Z1.ppp or Z4.aaa commits to the anchor cell fate while the other becomes another ventral uterine cell (Fig. 1) (Kimble, 1981; Kimble & Hirsh, 1979; Seydoux & Greenwald, 1989). We tested whether centralspindlin regulates this cell fate decision by analyzing somatic gonad precursor cells (SGPs) of *zen-4(RNAi)* and *cyk-4(RNAi)*-treated larvae expressing GFP-tagged HLH-2/E2A (Attner et al., 2019). HLH-2 is an E box transcription factor which drives expression of the Notch ligand *lag-2/Delta* during anchor cell specification (Karp & Greenwald, 2003, 2004). In addition to the distal tip cells, HLH-2 is expressed in the α and β cells in early L2 larva and by mid L2 expression is restricted to the AC. In empty vector RNAi controls, the four GFP::HLH-2-positive SGP nuclei were observed in early L2 larva before anchor cell specification (Fig. 3A). By mid L2, only the anchor cell nucleus continued to express GFP::HLH-2 (Fig. 3E). By comparison, *zen-4(RNAi)* treatment resulted in there being fewer GFP::HLH-2-positive SGP nuclei on average at early L2 (Fig. 3B,B’,D) with zero or two AC nuclei at mid L2 (Fig. 3F,F’,H). *cyk-4(RNAi)*-treated larvae exhibited similar phenotypes (Fig. 3C,C’,D,G,G’,H). Moreover, a somatic gonad cell nucleus reporter (Attner et al., 2019) was inspected at mid L2 which revealed *zen-4(RNAi)* and *cyk-4(RNAi)* larvae having significantly increased numbers of closely adjacent nuclei compared to empty vector (Fig. S2), consistent with cytokinesis defects. To directly determine if centralspindlin regulates the division of SGPs, we analyzed cells in a strain expressing membrane localized LIN-12/Notch and a LAG-2/Delta nuclear reporter (Medwig-Kinney et al., 2022) treated with *zen-4(RNAi)*. In the RNAi control, we observed anchor cells with a single nucleus positive for LAG-2 encircled by the LIN-12-expressing membrane at late L2 (Fig. 3I). As expected, *zen-4(RNAi)* treatment resulted in either the absence of cells expressing LAG-2 and LIN-12 (Fig. 3J,K), or the appearance of binucleate anchor cells with two LAG-2 positive nuclei inside a single LIN-12 expressing membrane (Fig. 3J’,K). In summary, centralspindlin promotes anchor cell specification through its regulation of SGP division. The variability of the phenotypes likely reflects random cytokinesis defects in the SGP lineage.

**Supplemental Figure S1.**
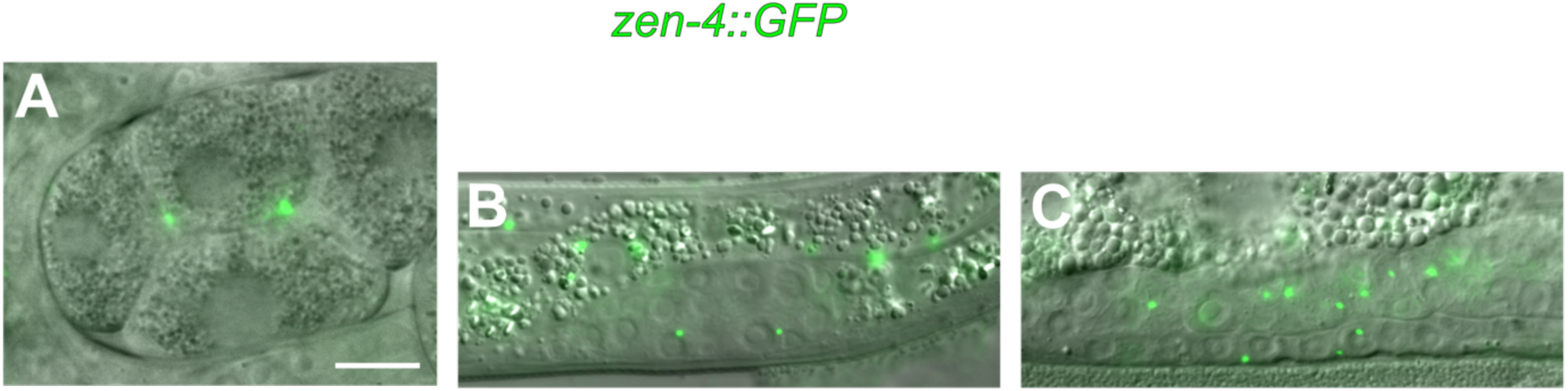
Centralspindlin subunit ZEN-4 is expressed during various stages of *C. elegans* development. ZEN-4::GFP-positive division remnants upon completion of cytokinesis can be seen in the developing embryo (A), early somatic gonad (B), vulval epithelium and overlying somatic gonad (C) of transgenic *zen-4::GFP* (*xsEx6*) expressed from the *zen-4* promoter. Scale bar, 10µm.

**Figure 3.**
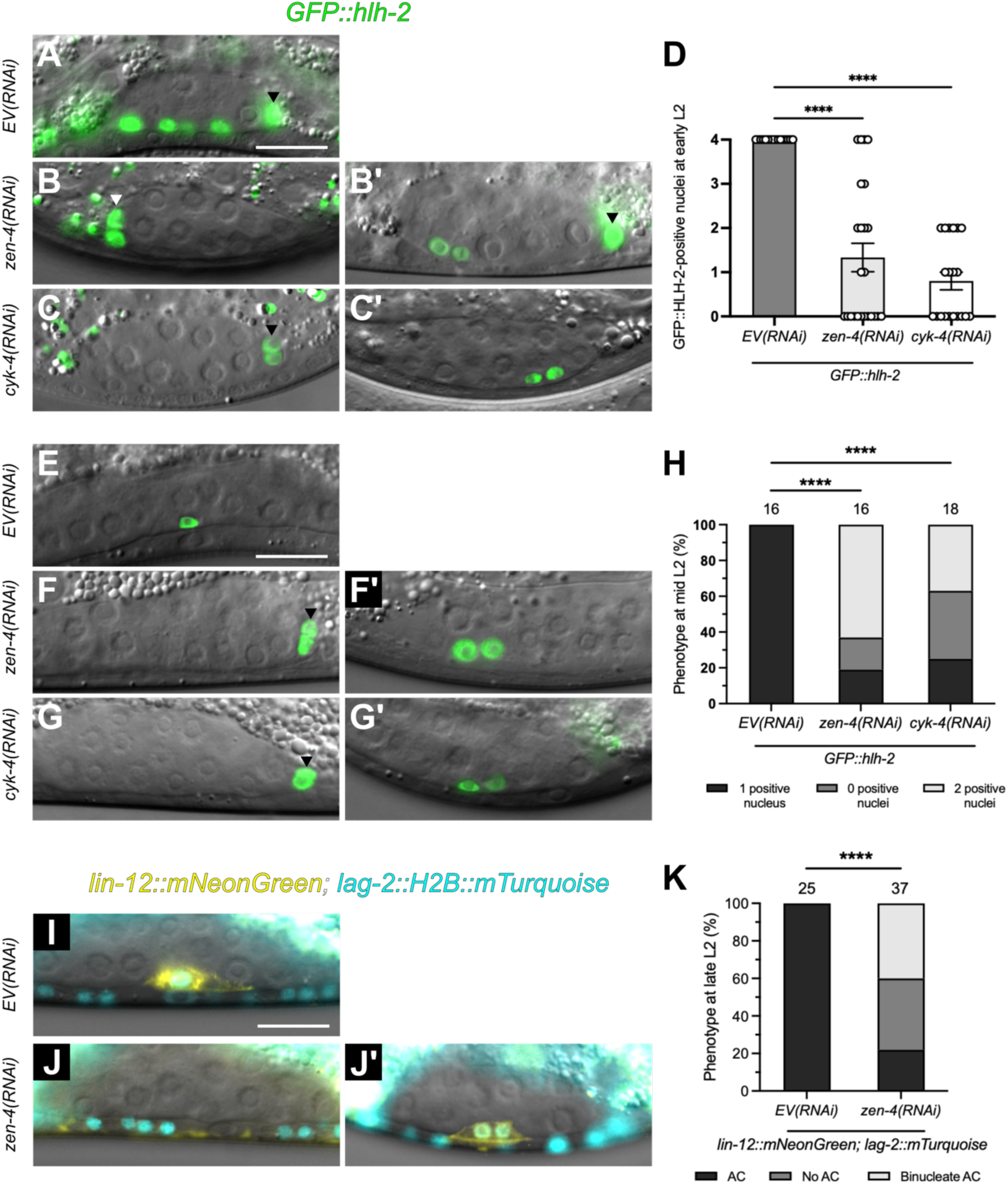
Centralspindlin promotes anchor cell specification by regulating somatic gonad precursor cytokinesis. (A-C’) Developing somatic gonads of AC specification reporter *GFP::hlh-2* (green) treated with *EV(RNAi)*, *zen-4(RNAi)* and *cyk-4(RNAi)* at early L2. (D) Quantification of the number of GFP-positive SGP nuclei in larvae treated with *EV(RNAi)*, *zen-4(RNAi)* and *cyk-4(RNAi)*. One-way ANOVA. \*\*\*\**P<0.*0001 (E-G’) *GFP::hlh-2* treated with *EV(RNAi)*, *zen-4(RNAi)* and *cyk-4(RNAi)* at mid L2. Arrowheads in (A,B’,C,F,G) denote distal tip cells that are also GFP::HLH-2 positive. Though not quantified these cells appear to be binucleate in larvae treated with *zen-4(RNAi)* and *cyk-4(RNAi)*. (H) AC specification defects in larvae treated with *EV(RNAi)*, *zen-4(RNAi)* and *cyk-4(RNAi)*. Chi-squared test with Bonferroni correction. \*\*\*\**P<0.0001.* (I-J’) LIN-12/Notch reporter *lin-12::mNeonGreen* (yellow) and LAG-2/Delta nuclear reporter *lag-2::H2B::mTurquoise* (cyan) expression at late L2 in *EV(RNAi)* and *zen-4(RNAi)* larvae. (K) AC specification defects in larvae treated with *EV(RNAi)* and *zen-4(RNAi)*. Chi-squared test with Bonferroni correction. \*\*\*\**P<0.0001.* Scale bars, 10µm.

**Supplemental Figure S2.**
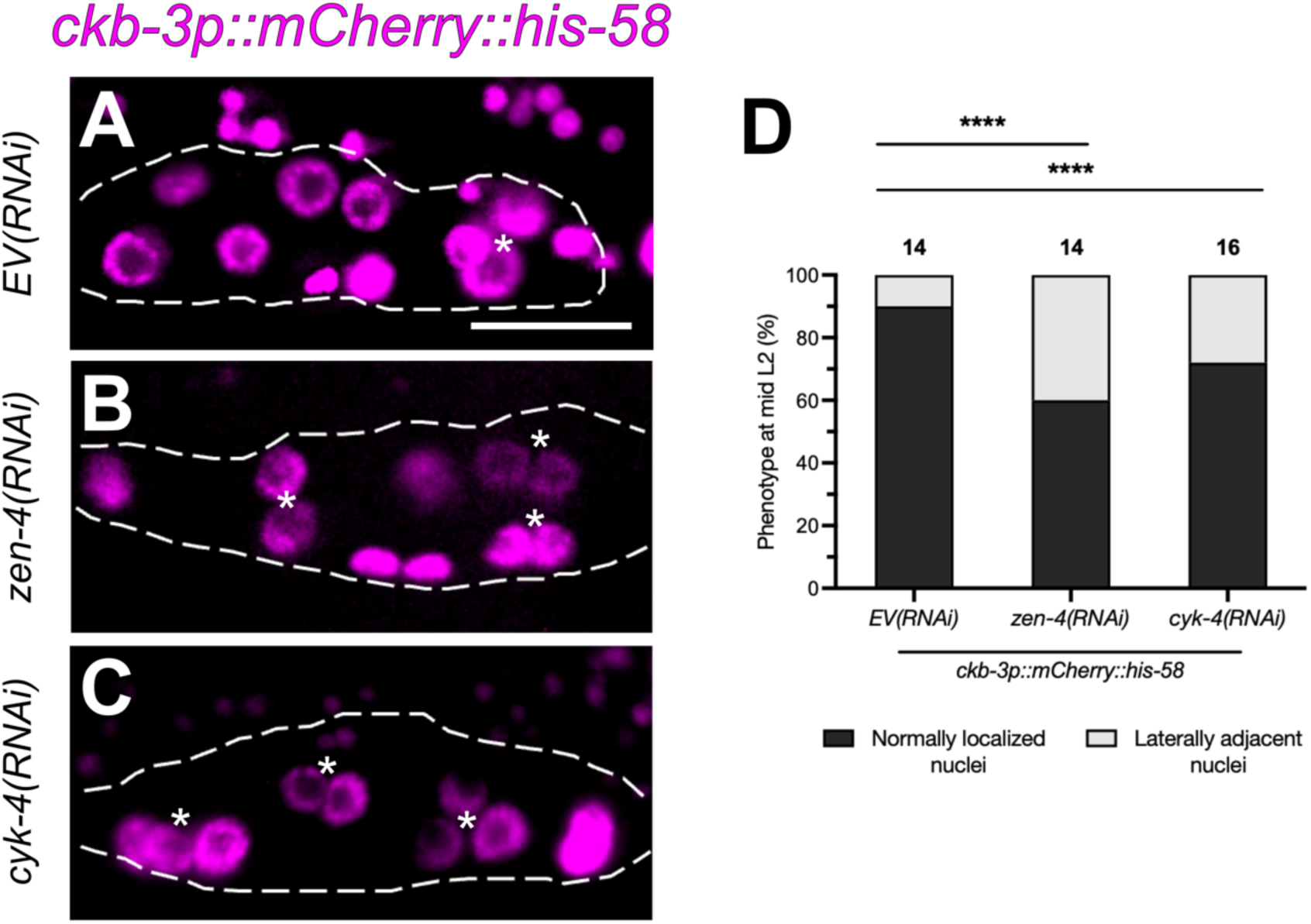
Knockdown of centralspindlin increases the occurrence of adjacent somatic gonad cell nuclei consistent with a cytokinesis defect. **(A-C)** Developing somatic gonads of mid L2 larvae expressing the somatic gonad cell nuclear reporter *ckb-3p::mCherry::his-58* (magenta) treated with *EV(RNAi)*, *zen-4(RNAi)* and *cyk-4(RNAi)*. Asterisks denote closely adjacent non-α cell nuclei that could result from a cytokinesis failure. (D) Characterization of nuclei in larvae treated with *EV(RNAi)*, *zen-4(RNAi)* and *cyk-4(RNAi)*. Chi-squared test with Bonferroni correction. \*\*\*\**P<*0.0001. Scale bars, 10µm.

### Binucleate anchor cells express LIN-3/EGF and can invade the basement membrane

The anchor cell performs multiple functions critical to vulval development (Gupta et al., 2012; Sternberg, 2005). We focused on two of these roles, in which the anchor cell secretes LIN-3/EGF required for VPC induction from early L3 (Hill & Sternberg, 1992), then invades the underlying basement membrane at late L3 to create the uterine-vulval connection necessary for egg laying (Sherwood & Sternberg, 2003). To determine whether centralspindlin regulates these functions, we analyzed a LIN-3/EGF reporter (Mereu et al., 2020) as well as a dual reporter of the anchor cell membrane and basement membrane (Hagedorn et al., 2013). In empty vector, we observed anchor cells with a single nucleus expressing LIN-3 at mid L3 (Fig. 4A), followed by formations of polarized invasive protrusions which successfully invaded basement membranes at late L3 (Fig. 4E). Consistent with the absence of an anchor cell, *zen-4(RNAi)* and *cyk-4(RNAi)* larvae often failed to express LIN-3 (Fig. 4B,C,D) and degrade the basement membrane (F,G,H,I). Interestingly, binucleate anchor cells observed in RNAi-treated larvae behaved similarly to anchor cells in empty vector in that they expressed LIN-3 (Fig. 4B’,C’,D) and invaded the basement membrane (Fig. 4F’,G’,H,I). They also appeared able to induce the VPCs; all *zen-4(RNAi)* larvae with nonzero VPC scores had a binucleate anchor cell (Fig. S3). These results demonstrate that centralspindlin is required for divisions that form the anchor cell but dispensable for LIN-3 secretion and basement membrane invasion.

**Figure 4.**
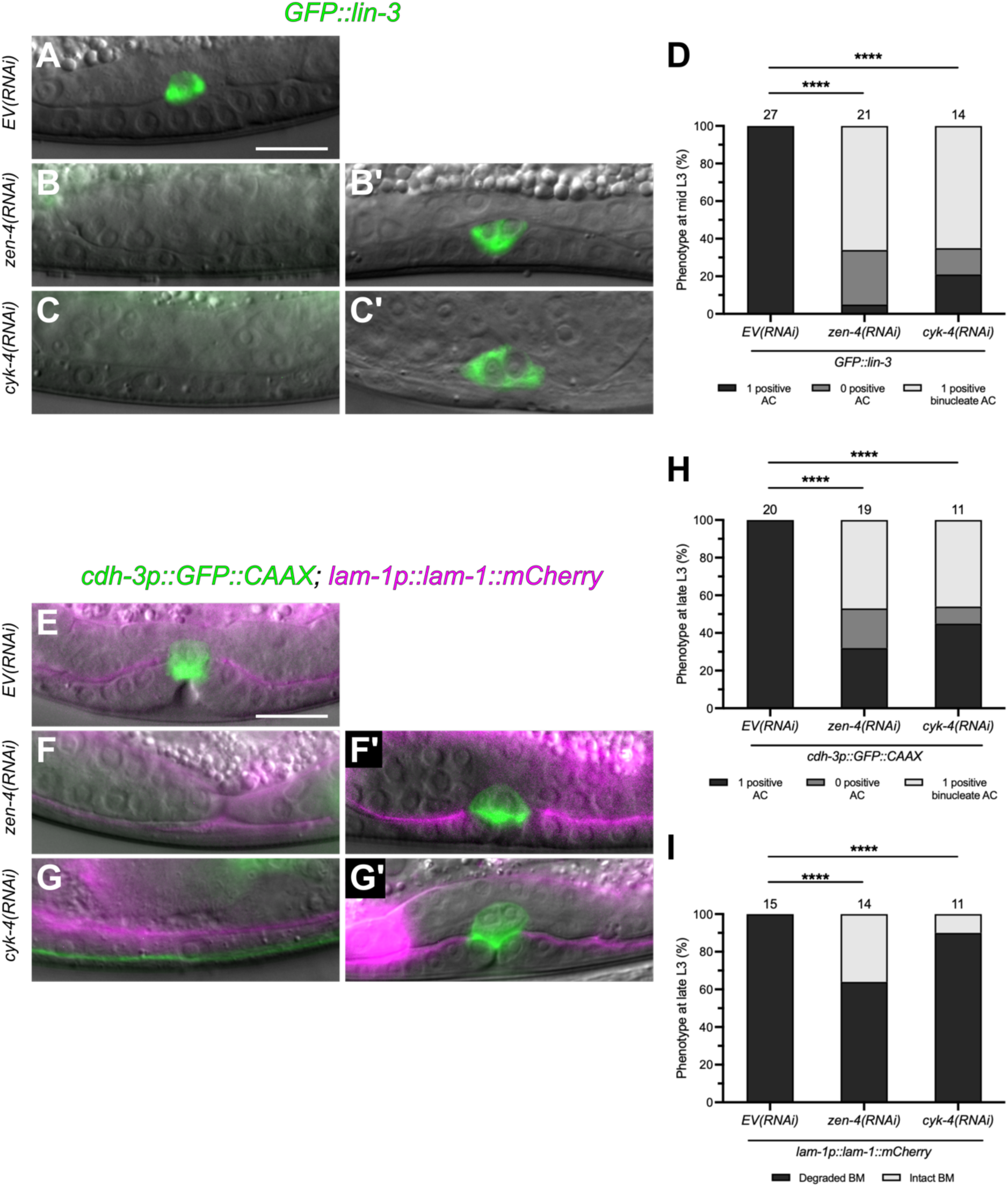
Binucleate anchor cells express LIN-3/EGF and invade the basement membrane. (A-C’) LIN-3/EGF reporter *GFP::lin-3* (green) treated with *EV(RNAi)*, *zen-4(RNAi)* and *cyk-4(RNAi)* at mid L2. (D) Quantification of LIN-3 expression defects in larvae treated with *EV(RNAi)*, *zen-4(RNAi)* and *cyk-4(RNAi)*. Chi-squared test with Bonferroni correction. \*\*\*\**P<0.0001.* (E-G’) Anchor cell membrane reporter *cdh-3p::GFP::CAAX* (green) and basement membrane reporter *lam-1p::lam-1::mCherry* (magenta) treated with *EV(RNAi)*, *zen-4(RNAi)* and *cyk-4(RNAi)* at late L2. *cdh-3p::GFP* is also expressed in the ventral nerve cord in (G). (H) Quantification of AC presence in larvae treated with *EV(RNAi)*, *zen-4(RNAi)* and *cyk-4(RNAi)*. Chi-squared test with Bonferroni correction. \*\*\*\**P<0.0001.* (I) Quantification of basement membrane degradation in larvae treated with *EV(RNAi)*, *zen-4(RNAi)* and *cyk-4(RNAi)*. Fisher’s exact test. Scale bars, 10µm.

**Supplemental Figure S3.**
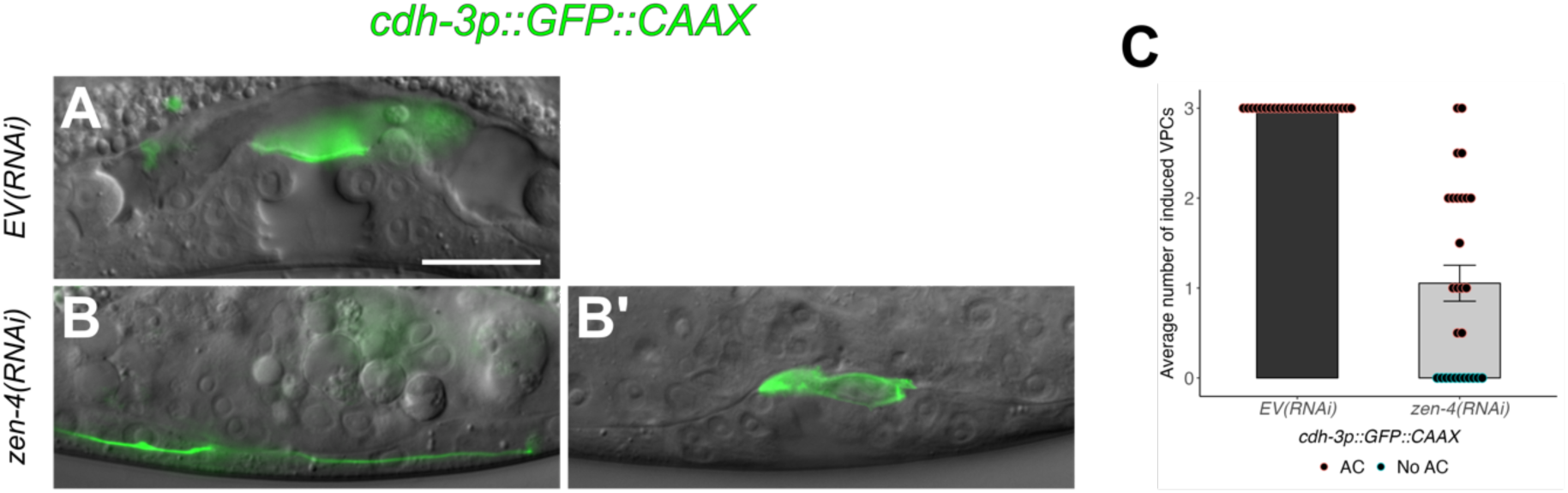
Binucleate anchor cells (ACs) induce the VPCs. (A-B’) AC membrane reporter *cdh-3p::GFP::CAAX* (green) treated with *EV(RNAi)* and *zen-4(RNAi)* at mid L4. (B) Absence of the AC results in zero vulva induction. *cdh-3p::GFP* is also expressed in the ventral nerve cord. (B’) Presence of the AC correlates with partial vulva induction. (C) Quantification of VPC induction of empty vector and *zen-4(RNAi)* in relation to absence or presence of the AC. Scale bars, 10µm.

### ZEN-4 is primarily required during AC specification to regulate vulva induction

To test if centralspindlin is required at the time of anchor cell specification to regulate vulva induction, we performed temperature shifts of *zen-4(or153)* and *cyk-4(t1689)* from the permissive temperature of 15℃ to the nonpermissive temperature of 26℃ to inactivate centralspindlin either during anchor cell specification (during L2) or VPC induction (from L3). Shifting *zen-4(or153)* and *cyk-4(t1689)* larvae to 26℃ during L2 was sufficient to induce a strong vulvaless phenotype similar to larvae shifted from L1 (Fig. 5A,B,D,E) while larvae shifted from L3 had greater average numbers of induced VPCs (Fig. 5C,F). Importantly, all larvae shifted from L3 had nonzero VPC scores (Fig. 5G). These results show that centralspindlin promotes vulval development primarily by regulating anchor cell specification which indirectly affects VPC induction.

**Figure 5.**
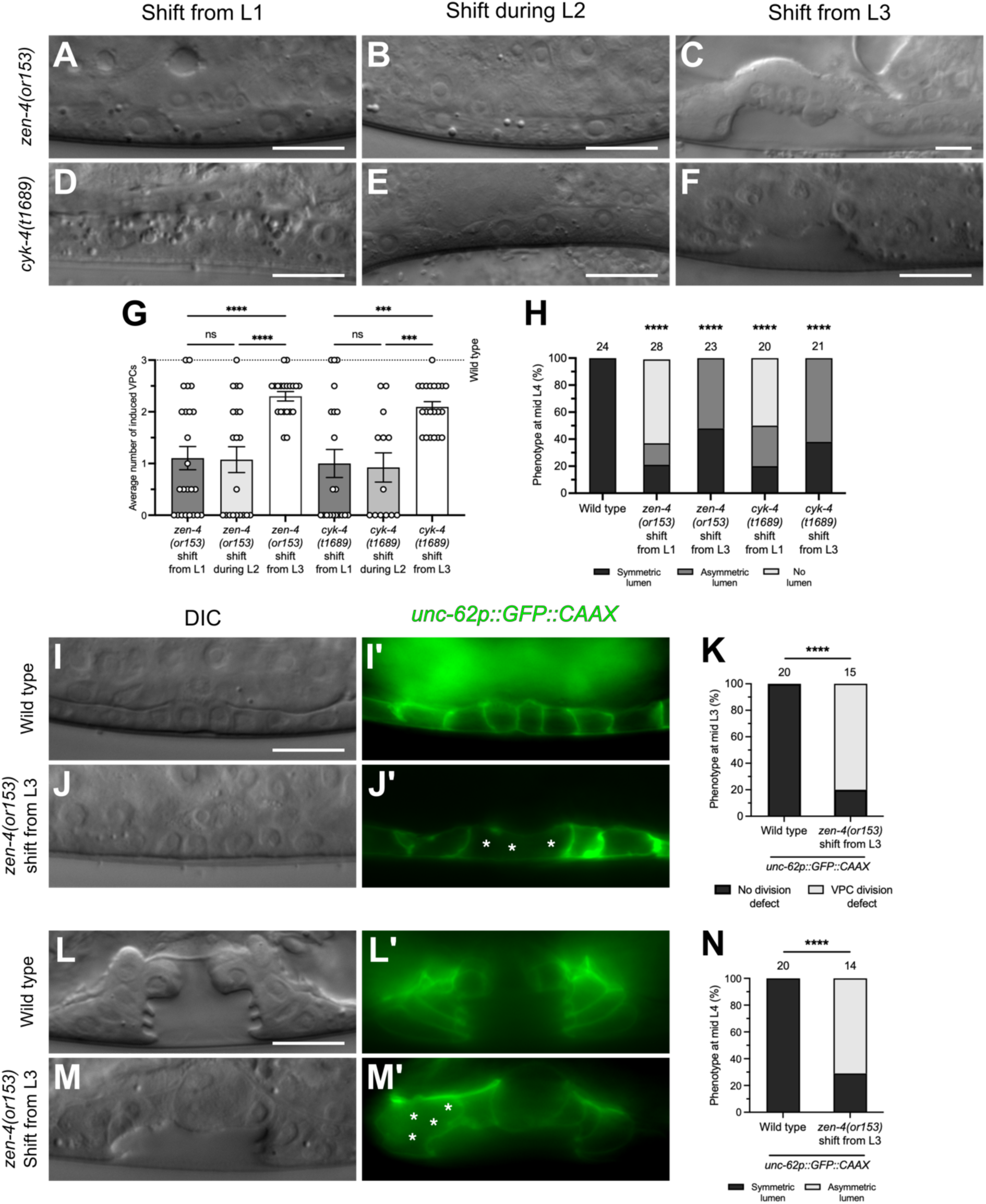
Centralspindlin regulates VPC division and vulval morphogenesis. (A-F) Developing vulva of mid L4 stage *zen-4(or153)* and *cyk-4(t1689)* larvae grown at restrictive temperature from L1, during L2, and from L3. (G) Quantification of VPC induction of *zen-4(or153)* larvae grown at restrictive temperature from L1, during L2, and from L3. One-way ANOVA. ns, not significant, \*\*\*\**P<0.0001.* (H) Lumen morphology in wild type, *zen-4(or153)* and *cyk-4(t1689)* in restrictive temperature from L1 and from L3. Chi-squared test with Bonferroni correction. \*\*\*\**P<0.0001.* (I-J’) Differential interference contrast (DIC) and epifluorescence micrographs of VPC membrane reporter *unc-62p::GFP::CAAX* (green) and *zen-4(or153); unc-62p::GFP::CAAX* in restrictive temperature from L3 at mid L3. (K) Quantification of VPC division defects in *unc-62p::GFP::CAAX* and *zen-4(or153); unc-62p::GFP::CAAX* in restrictive temperature from L3. Fisher’s exact test. \*\*\*\**P<0.0001.* (L-M’) Developing vulva of *unc-62p::GFP::CAAX* and *zen-4(or153); unc-62p::GFP::CAAX* in restrictive temperature from L3 at mid L4. Asterisks in (J’,M’) denote nuclei of multinucleate vulval cells in (J,M). (N) Quantification of lumen formation defects in *unc-62p::GFP::CAAX* and *zen-4(or153); unc-62p::GFP::CAAX* in restrictive temperature from L3. Fisher’s exact test. \*\*\*\**P<0.0001.* Scale bars, 10µm.

### ZEN-4 is required for vulval cell cytokinesis and vulva morphology

Although maintaining *zen-4(or153)* and *cyk-4(t1689)* larvae in permissive temperature until L3 restored VPC induction, it also led to vulval morphogenesis defects in the form of asymmetric lumens (Fig. 5H). Notably, we often observed extended invaginations in which VPC daughters appeared to have divided longitudinally (Fig. 5C,F). A similar phenotype was reported to be associated with defects in VPC division orientation and migration (Kishore & Sundaram, 2002). To determine if this abnormal morphology could be attributed to cytokinesis defects in the vulval cells, we analyzed *zen-4(or153)* larvae expressing a GFP::CAAX membrane marker in the VPCs (Matus et al., 2014). We found that most *zen-4(or153)* larvae shifted to restrictive temperature from L3 had multinucleate VPC daughter cells at mid L3 (Fig. 5J,J’,K), and these division defects correlated with lumen morphology defects at mid L4 (Fig. 5M,M’,N). However, we also observed that all *zen-4(or153)* larvae upshifted from L3 exhibited some degree of vulva induction (Fig. 5C,G,H,M), consistent with centralspindlin function during anchor cell fate specification being sufficient for vulva induction. Therefore, during L3 stage, centralspindlin promotes vulva cell cytokinesis and vulva morphogenesis.

### ZEN-4 is required in the somatic gonad for vulva induction

To further characterize the tissue specific requirements for centralspindlin during vulval development, we performed rescue of *zen-4(or153)* through transgenic expression of ZEN-4::mCherry driven by either the *lin-31* promoter specific to the VPCs (Tan et al., 1998) or *ckb-3* promoter specific to the SGPs (Kroetz & Zarkower, 2015) (Fig. 6A,B). VPC-specific *zen-4* expression did not restore vulval induction (Fig. 6C). However, vulval morphology was largely restored in larvae which underwent VPC induction and expressed the transgene (Fig. 6A’). In contrast, SGP-specific rescue led to increased VPC induction scores and lumen formation, albeit with morphology defects (Fig. 6B’,C).

**Figure 6.**
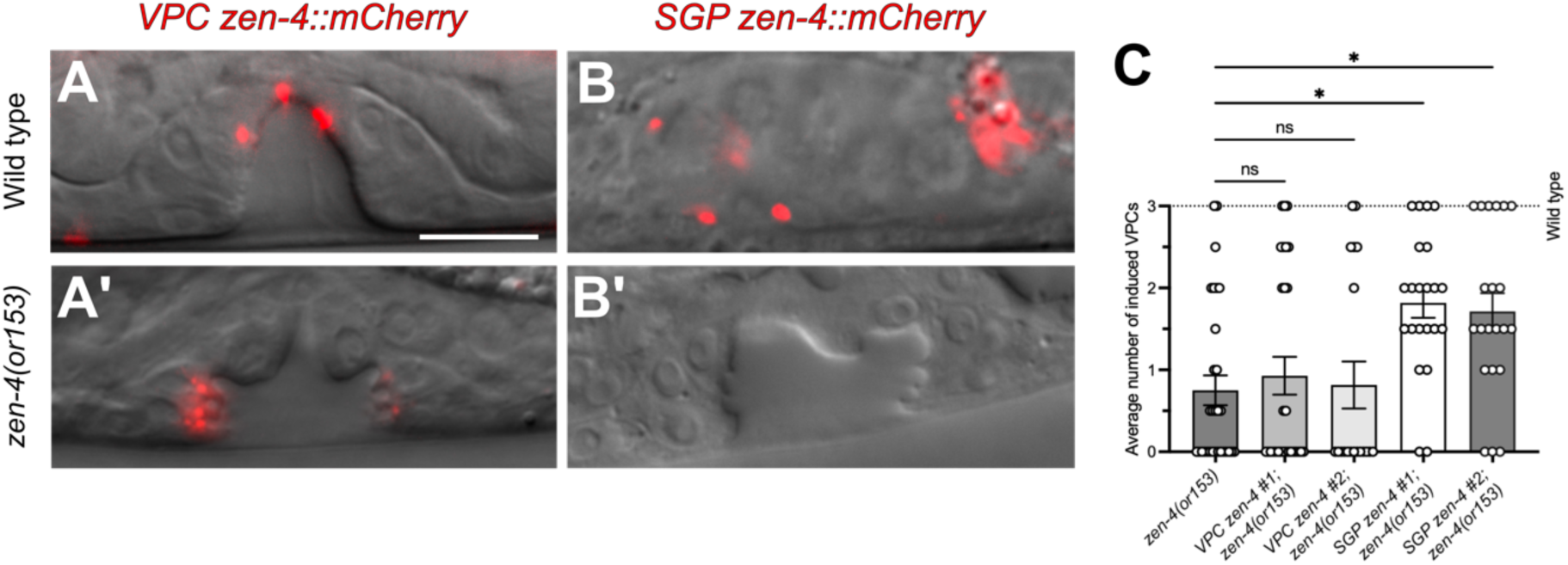
Somatic gonad precursor cell (SGP)-specific *zen-4* expression rescues VPC induction. (A) Developing vulva of wild type expressing vulval precursor cell (VPC)-specific *zen-4* (red) at early L4. (A’) Developing vulva of *zen-4(or153)* expressing VPC-specific *zen-4* at mid L4. (B) Developing wild type somatic gonad expressing SGP-specific *zen-4* at early L2. (B’) Developing vulva of *zen-4(or153)* expressing SGP-specific *zen-4* at mid L4. (C) Quantification of VPC induction of *zen-4(or153)*, *VPC zen-4; zen-4(or153)*, and *SGP zen-4; zen-4(or153)*. Two independent transgenic lines (#1 and #2) were analyzed. One-way ANOVA. ns, not significant. \**P<0.05.* Scale bars, 10µm.

## DISCUSSION

We demonstrate that centralspindlin is required for vulval development by regulating cytokinesis in both the somatic gonad and the vulval cells. In the somatic gonad, centralspindlin regulates SGP cytokinesis necessary for anchor cell specification which is critical for VPC induction. Centralspindlin also functions in the underlying vulval epithelium to promote vulva morphogenesis by regulating the divisions of the VPCs and their descendants. While centralspindlin function is essential for completion of cytokinesis during early embryogenesis, we find that centralspindlin has different threshold requirements for postembryonic divisions in the somatic gonad and vulva. Similarly, the CYK-4 GAP domain, while essential during embryogenesis, appears to not be required for cytokinesis during vulva development. When specified, the binucleate anchor cell is largely functional in its ability to signal and invade the basement membrane.

Centralspindlin is indirectly required for vulva induction. We originally hypothesized that centralspindlin would be required in the VPCs for vulva induction. However, *zen-4(RNAi)* failed to suppress the multivulva phenotypes of *lin-1* and *lin-15a/b* mutants, suggesting that ZEN-4 functions upstream of the LET-23 EGFR signaling pathway to regulate vulva induction. One caveat of this experiment is that RNAi is less efficient in the VPCs (Bourdages et al., 2014; Matus et al., 2014). However, the fact that the vulvaless phenotype of *zen-4(RNAi)* is as severe as that of the *zen-4(or153)* mutant supports an upstream role for ZEN-4 in the somatic gonad.

Centralspindlin regulates cytokinesis of the somatic gonad precursor cells for AC specification and vulva induction. The AC fate is specified during the L2 stage. Using markers for the somatic gonad precursor cells and the AC we demonstrate that centralspindlin regulates cytokinesis of the AC precursors. As a result, when centralspindlin is reduced the AC is often not specified or binucleate when present. The presence of the AC, binucleate or not, correlates with vulva induction, indicating that centralspindlin’s role in vulva induction is due to its function in the somatic gonad to specify the AC fate. Consistent with this, *zen-4(or153)* and *cyk-4(t1689)* temperature shift experiments demonstrate that ZEN-4 and CYK-4 are required for vulva induction during the L2 stage when the AC is specified. Shifting *zen-4(or153)* or *cyk-4(t1689)* to non-permissive temperature during the L3 stage when AC to VPC signaling occurs has little effect on vulva induction. Moreover, expression of ZEN-4 in the somatic gonad, but not the VPCs, is sufficient to rescue the vulva induction phenotype of *zen-4(or153).* Thus, ZEN-4 functions in the somatic gonad to promote AC specification and subsequent vulva induction.

Centralspindlin regulates vulva morphogenesis. While incomplete vulva induction results in an abnormal vulva, *zen-4(or153)* or *cyk-4(t1689)* temperature shift to non-permissive temperature during the L3 stage reveals that centralspindlin is required for vulva morphogenesis. In addition, expression of ZEN-4 in the VPCs rescues the morphogenesis phenotype of *zen-4(or153)*. Using a membrane marker expressed in the VPCs, it is apparent that centralspindlin is required for completion of cytokinesis in the VPC daughters. Although cytokinesis defects are most likely contributing to the morphogenesis phenotype, we cannot rule out other potential functions by centralspindlin that affect vulva morphogenesis. The vulva morphogenesis phenotype shows some similarity to those described for larvae with mutations of the *ced-10* and *mig-2* Rac GTPases (Kishore & Sundaram, 2002) in that the vulval lumen is enlarged, thus it is possible that centralspindlin contributes to vulva morphogenesis via regulation of cell polarity or cell migration. Furthermore, in the VPC rescue experiments we note that ZEN-4::mCherry midbody remnants localize to the luminal apical membrane of the vulva as they do in embryonic epithelial cells (Bai et al., 2020), potentially acting as polarity cues.

Although the *cyk-4(t1689)* coiled coil mutant and *cyk-4(RNAi)* exhibited defects in vulva induction and anchor cell specification, we did not observe any abnormal vulval developmental phenotypes in *cyk-4(or749)* mutants. The E448K substitution in *cyk-4(or749)* compromises its GAP domain at restrictive temperatures and inhibits embryonic cytokinesis (Canman et al., 2008). Our results suggest that CYK-4’s GAP activity is not essential for postembryonic cell proliferation in the somatic gonad and vulval epithelium and thus may not function as a GAP regulating CED-10 and MIG-2 Rac GTPases during vulva morphogenesis. Our findings corroborate reports of *cyk-4(or749)* not affecting larval growth (Lee et al., 2018) or spermatheca morphogenesis (Zhang et al., 2023), both of which are affected by *zen-4* and *cyk-4* depletion. We speculate that CYK-4 primarily contributes to vulval development by interacting through its coiled coil domain with ZEN-4 which is required for complex assembly (Mishima et al., 2002) and midzone localization (Jantsch-Plunger et al., 2000).

The requirements for centralspindlin when completing cytokinesis are reduced during postembryonic divisions of the SGP and VPCs. The *zen-4(or153)* mutation is a fast-acting temperature sensitive mutant (Severson et al., 2000); shifting to non-permissive temperature results in mislocalization of CYK-4 from the central spindle in minutes with subsequent failure to complete cytokinesis (Jantsch-Plunger et al., 2000). *zen-4(or153)* and *cyk-4(ts1689)* at non-permissive temperature result in 100% penetrant embryonic lethality due to multinucleate embryos that fail to complete cytokinesis. We find that shifting L1 larvae to non-permissive temperature largely results in larval lethality, but there are animals which survive to the L4 stage, indicating that cytokinesis is occurring. This is evident in the VPCs and their descendants in which some cells complete cytokinesis despite the absence of centralspindlin function, as assessed with a VPC membrane marker. In daughter cells with completed cytokinesis, centralspindlin may not be essential or may have lower requirement thresholds such that residual amounts are sufficient to complete cytokinesis. Such variability may be due to tissue-specific differences in factors such as cell size, cell polarity, microtubule abundance or cell-cell interactions. Consistent with this idea, in *spd-1* mutants cytokinesis completes during early embryonic divisions but fails further down the lineage such as that of the EMS cell (Verbrugghe & White, 2004).

LIN-12/Notch and LIN-3/EGF signaling are largely functional in binucleate VU and AC cells. LIN-12/Notch signaling between Z1.ppp and Z4.aaa determines which cell adopts the VU and AC fate. When centralspindlin is reduced the AC is typically binucleate or not specified, but we do not observe tetranucleate ACs. Thus, LIN-12/Notch signaling most likely occurs normally between binucleate Z1.pp and Z4.aa cells during specification of a binucleate AC, whereby animals with no AC may reflect cytokinesis defects which occurred earlier in the Z1 and Z4 lineages that negate LIN-12/Notch signaling since LIN-12 expression is only visible after the second divisions (Wilkinson & Greenwald, 1995). When specified, the binucleate AC appears to be largely functional in that it expresses the LIN-3/EGF ligand. Furthermore, its presence correlates with vulva induction and basement membrane degradation. However, vulva induction is not all or nothing in centralspindlin mutants; we frequently see partial inductions which also correlates with binucleate AC presence. This suggests that the binucleate AC could have reduced LIN-3/EGF signaling or its underlying VPCs have reduced potential to respond to the inductive signal. Nevertheless, our results are consistent with previous studies demonstrating that multinucleate cells can maintain functionality in the absence of centralspindlin (Hardin et al., 2008).

In conclusion, centralspindlin controls vulval development by ensuring the fidelity of cytokinesis in the somatic gonad and vulval precursor cells. Our results further demonstrate how cell division and cell fate specification need to be coordinated during animal development.

## MATERIALS AND METHODS

### Strains and maintenance

*C. elegans* strains were maintained on nematode growth medium agar plates seeded with HB101 *E. coli* (Stiernagle, 2006). Strains were maintained at 20℃ apart from temperature sensitive strains that were maintained at 15℃ or 26℃ where indicated. The Bristol strain N2 was used as wild type control. Genetic crosses and the generation of new strains were done as previously described (Brenner, 1974). Transgenic strains for tissue specific expression of *zen-4* were generated by DNA microinjection of plasmids containing *lin-31p::mCherry::zen-4* or *ckb-3p::mCherry::zen-4* (40ng/µl) with the *ttx-3p::GFP* (80ng/µl) or *myo-2p::mCherry* (40ng/µl) co-injection marker as described (Berkowitz et al., 2008).The strains used in this study are listed in Supplementary Table S1.

### Plasmid construction

To construct *lin-31p::mCherry::zen-4*, the *zen-4* ORF was amplified from a plasmid containing the M03D4 cosmid (Kaitna et al., 2000) using the primers 5’-CAATGGTACCTCGTCGCGTAAACGAG-3’ (forward) and 5’-CAATGAGCTCTTACTTTCGTGTGCTCGGAG-3’ (reverse). The PCR product was digested with KpnI and SacI, then inserted into a plasmid expressing the *lin-31* promoter and mCherry (Gauthier & Rocheleau, 2021). To construct *ckb-3p::mCherry::zen-4*, the *ckb-3* promoter was amplified from the pPD95.75-*ckb-3Prom* plasmid (Kroetz & Zarkower, 2015) using the primers 5’-AAGAAAGGCACCGTCCAATCATCGACGCCGGCGTTTTTGCTCACCTAAAAAAAATACC-3’ (forward) and 5’-CCTTTGGGTCCTTTGGCCAATCCCGGGGATCCGTTAATTTAGCAGCTTTTGAGAAATG-3’ (reverse). *mCherry::zen-4* was digested from *lin-31p::mCherry::zen-4* with NaeI and BamHI, then assembled with the *ckb-3* promoter via Gibson assembly (New England Biolabs, US).

### Temperature shift experiments

Temperature sensitive strains were synchronized as described (Stiernagle, 2006) at 15℃, then temperature shifts were performed based on previously reported growth times (Byerly et al., 1976). For positive controls, synchronized L1 larvae were incubated at 26℃ for 36 hours to be imaged at mid L4. For shifts during L2, synchronized L1 larvae were incubated at 15℃ for 18 hours, then at 26℃ for 10 hours, then at 15℃ for 20 hours to be imaged at mid L4. For shifts from L3, synchronized L1 larvae were incubated at 15℃ for 32 hours, then at 26℃ for 11 hours to be imaged at mid L4.

### RNA-mediated interference

RNA-mediated interference (RNAi) by feeding was performed as described (Kamath et al., 2001) using the empty vector (L4440) as a negative control, *zen-4* (IV-3I07), *cyk-4* (III-6H12), and *air-2* (I-9A05) clones of the Ahringer RNAi Library (Source BioScience, UK). To avoid embryonic lethality, L1 larvae were synchronized as described (Stiernagle, 2006), transferred onto seeded RNAi plates and imaged at various developmental stages. RNAi clones were verified by DNA sequencing.

### Microscopy

Larvae were transferred to a 2% agarose pad in 10mM levamisole on a slide with a coverslip of 0.13 to 0.16mm thickness. Differential interference contrast and epifluorescence images were acquired on an Axio Imager A1 using the Axiocam 305 mono or AxioCam mRm camera with AxioVision software (Zeiss, Germany).

### VPC induction scoring

Quantification of VPC induction was performed as described (Gauthier & Rocheleau, 2017). Briefly, the VPCs of mid L4 stage larvae were scored 0, 0.5 or 1 based on the number of descendent nuclei counted. Larvae with total scores of 3 were classified as wild type, those with less than 3 as vulvaless, and those with more than 3 as multivulva.

### Statistical analysis

Statistical analysis and graphs were primarily done using Graphpad Prism version 10 as well as R for Supplementary Figure 3.

### Resources

Data deposited in Wormbase (wormbase.org) was essential in the planning and design of the research described (Sternberg et al., 2024).

## Abbreviations

AC: (anchor cell)
EGFR: (epidermal growth factor receptor)
RNAi: (RNA-mediated interference)
SGP: (somatic gonad precursor cell)
VPC: (vulval precursor cell)

## ACKNOWLEDGEMENTS

We are indebted to Jung Hwa Seo for her technical guidance and for generating the QR1086 and QR1087 tissue-specific *zen-4* transgenic strains. We thank Alisa Piekny (Concordia University) for critically reading the manuscript. We thank the Piekny (Concordia University), Greenwald (Columbia University), Glotzer (University of Chicago), Hajnal (University of Zurich), Kroetz (Bellarmine University), Matus (Stony Brook University), Pani (University of Virginia), Sherwood (Duke University) and Van Raamsdonk (McGill University) labs for their kind gifts of strains and reagents. Some strains were provided by the Caenorhabditis Genetics Center (CGC), which is funded by NIH Office of Research Infrastructure Programs (P40 OD010440). Confocal microscopy was conducted in the Molecular Imaging Platform of the RI-MUHC who is supported in part by the Fonds de recherche du Québec – Santé (FRQS). This work was funded by a Natural Sciences and Engineering Research Council of Canada (NSERC) Discovery Grant (RGPIN-2018-05673) to CER.

---

**Supplementary Table S1.**
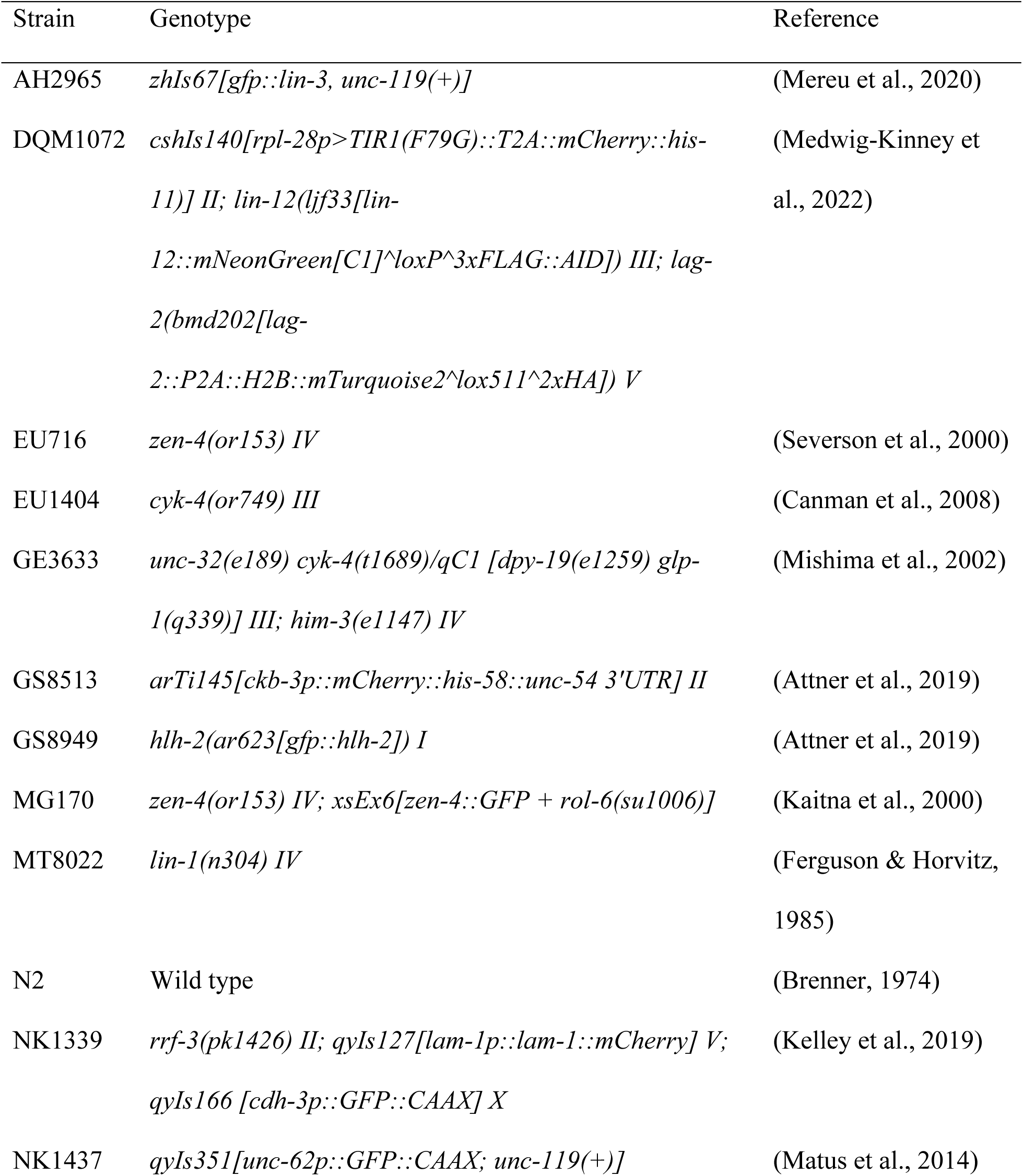
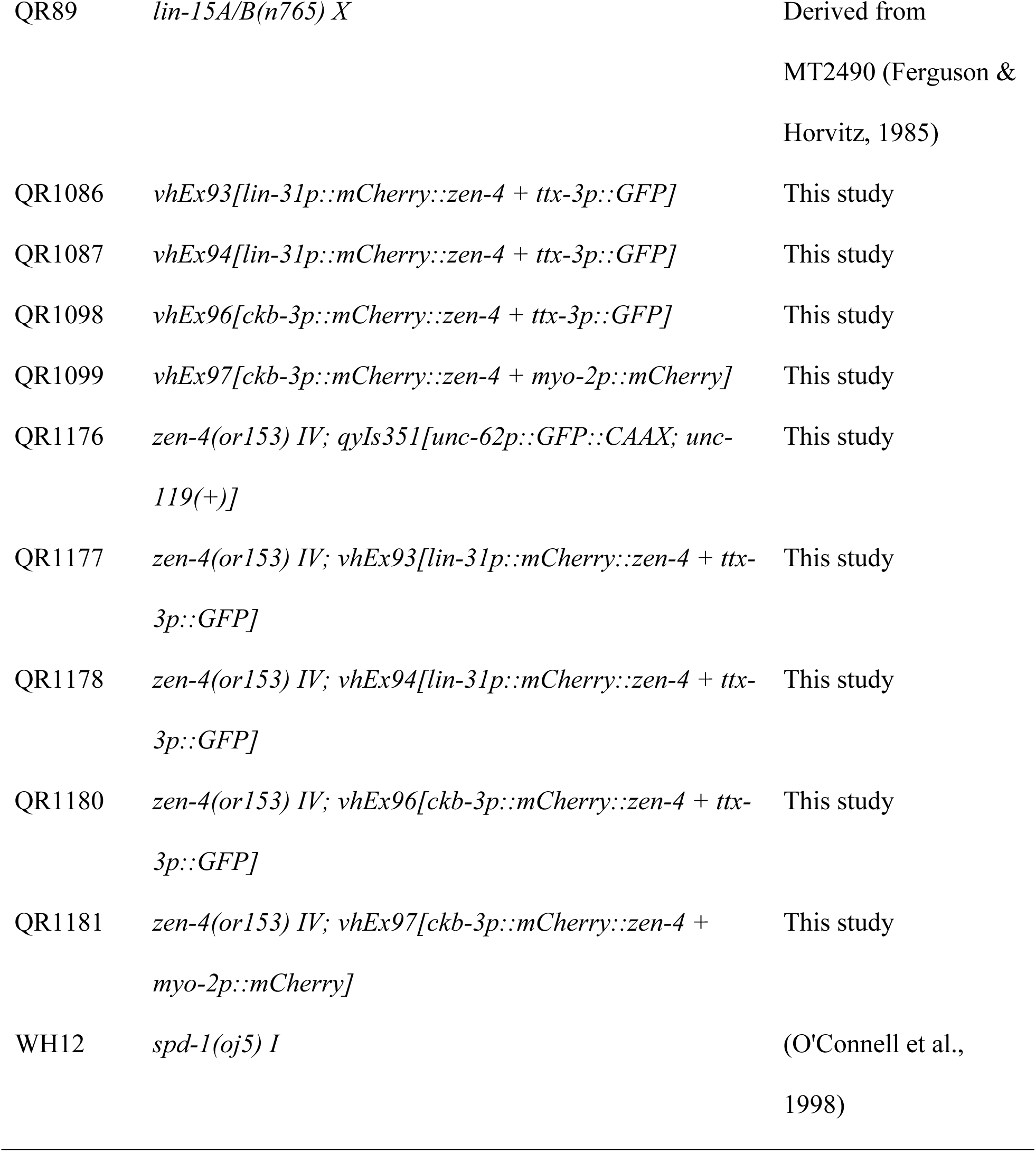

